# A high-throughput screen for transcription activation domains reveals their sequence characteristics and permits reliable prediction by deep learning

**DOI:** 10.1101/2019.12.11.872986

**Authors:** Ariel Erijman, Lukasz Kozlowski, Salma Sohrabi-Jahromi, James Fishburn, Linda Warfield, Jacob Schreiber, William S. Noble, Johannes Söding, Steven Hahn

## Abstract

Transcription activation domains (ADs) are encoded by a wide range of seemingly unrelated amino acid sequences, making it difficult to recognize features that permit their dynamic behavior, fuzzy interactions and target specificity. We screened a large set of random 30-mer peptides for AD function and trained a deep neural network (ADpred) on the AD-positive and negative sequences. ADpred correctly identifies known ADs within protein sequences and accurately predicts the consequences of mutations. We show that functional ADs are (1) located within intrinsically disordered regions with biased amino acid composition, (2) contain clusters of hydrophobic residues near acidic side chains, (3) are enriched or depleted for particular dipeptide sequences, and (4) have higher helical propensity than surrounding regions. Taken together, our findings fit the model of “fuzzy” binding through hydrophobic protein-protein interfaces, where activator-coactivator binding takes place in a dynamic hydrophobic environment rather than through combinations of sequence-specific interactions.

## Introduction

Transcription activators are regulatory factors that stimulate transcription in response to signaling pathways controlling processes such as development, growth and stress response (Levine et al., 2014; Spitz and Furlong, 2012). Misregulation of activators or mutations within them can lead to many human diseases and syndromes (Bradner et al., 2017). Each activator contains one or more transcription activation domains (ADs) that usually target coactivators – complexes that contact the basal transcription machinery and/or have chromatin modifying activity (Erkina et al., 2016; Hahn and Young, 2011). AD-coactivator binding initiates a series of events leading to productive transcription initiation, elongation and reinitiation. There are hundreds of cellular activators with distinct ADs, but most target only a small number of coactivators such as Mediator, TFIID, Swi/Snf, SAGA, NuA4 and p300. Broadly acting ADs can target several of these coactivators, allowing them to act on a large set of genes with different coactivator requirements. ADs have also been implicated in promoting the formation of intracellular condensates at enhancers, triggering the recruitment of a large dynamic network of coactivators and other factors responsible for gene activation (Boija et al., 2018; Cho et al., 2018; Chong et al., 2018; Shrinivas et al., 2019).

Nearly all characterized eukaryotic ADs are contained within intrinsically disordered protein regions (IDRs) that lack a stable 3D structure in the absence of a binding partner (Brzovic et al., 2011; Currie et al., 2017; Kussie et al., 1996; Sugase et al., 2007; Uesugi et al., 1997). In many systems apart from transcription, molecular recognition by IDRs is mediated by short linear motifs (SLMs), 3-10 residue sequence motifs found in otherwise unrelated proteins (Das et al., 2012; Nguyen Ba et al., 2012). In contrast, AD function is encoded by a wide range of seemingly unrelated sequences. For example, while AD sequences can be moderately conserved in closely related orthologs (Pacheco et al., 2018), no common sequence motif has been found when comparing ADs from different transcription factors. In addition, small-scale screens for ADs using random sequences of varying length fused to a DNA binding domain found that ∼1% of these sequences encoded AD function (Abedi et al., 2001; Erkine et al., 2002; Ma and Ptashne, 1987a; Ravarani et al., 2018; Ruden et al., 1991). Finally, prior work analyzing natural and synthetic ADs have found several sequence features that correlate with AD function including intrinsic disorder, the presence of acidic, hydrophobic, and aromatic residues, low sequence complexity, net negative charge (or lack of positive charge) and, in some cases, alpha helix propensity (Brzovic et al., 2011; Dyson and Wright, 2016; Erkina et al., 2016; Ma and Ptashne, 1987a; Ravarani et al., 2018).

Recent structural work and molecular analysis suggests that a large class of ADs recognize coactivators via a dynamic “fuzzy” protein-protein interface. For example, the yeast activator Gcn4 contains tandem ADs that bind four structured domains in the Mediator subunit Med15 (Brzovic et al., 2011; Tuttle et al., 2018; Warfield et al., 2014). Structural analysis showed that the individual AD-Med15 interactions are dynamic, and that the two factors appear to interact via a cloud of hydrophobicity rather than through sequence-specific interactions. This binding mechanism does not require a unique sequence motif for AD function. Because of this, it has been difficult to predict sequences with AD function and to understand which features promote their dynamic binding properties and specificity. Understanding these fundamental properties of ADs is essential toward progress in determining the molecular basis of AD specificity for certain coactivators, dissecting mechanisms used in gene activation, and in predicting the consequences of naturally occurring mutations on AD function.

In this work, we used a high throughput approach to screen over a million synthetic peptide sequences and found large numbers of AD-positive and AD-negative sequences. We analyzed the resulting sequence sets using logistic regression and also developed a deep neural network (NN) predictor of AD function, termed ADpred. The combination of these two approaches allowed us to identify sequence features that specify AD function. ADpred also enables identification of peptides having AD function within the yeast proteome and accurately predicts the consequences of mutations on AD function.

## Results

### A high-throughput screen for synthetic activation domains

To identify features encoding AD function, we isolated many synthetic ADs using a high throughput approach. We reasoned that gathering large sets of polypeptides with and without AD function would allow computational identification of physical properties, sequence motifs, and other features associated with ADs. Well-characterized natural ADs range from ∼5 to > 100 residues in length, but many are shorter than 30 residues. To generate large numbers of synthetic ADs, we designed libraries that encode 30 randomized amino acids attached to the N-terminal linker region and DNA-binding domain of Gcn4 (residues 132-281) (**Fig 1A**). Prior work showed that this Gcn4 derivative has no inherent AD function and that it can accept a wide variety of natural and synthetic ADs, permitting activation of yeast Gcn4-dependent genes (Pacheco et al., 2018; Warfield et al., 2014). By varying the ratio of the four DNA bases separately at codon positions 1, 2, and 3 (Labean and Kauffman, 1993), we made two libraries that either (1) biased the randomized coding sequences for residues normally enriched in IDRs (Uversky, 2014) or (2) that encoded an approximately equal representation of all amino acids. The results reported below are derived from analysis of the combined libraries.

**Figure 1.**
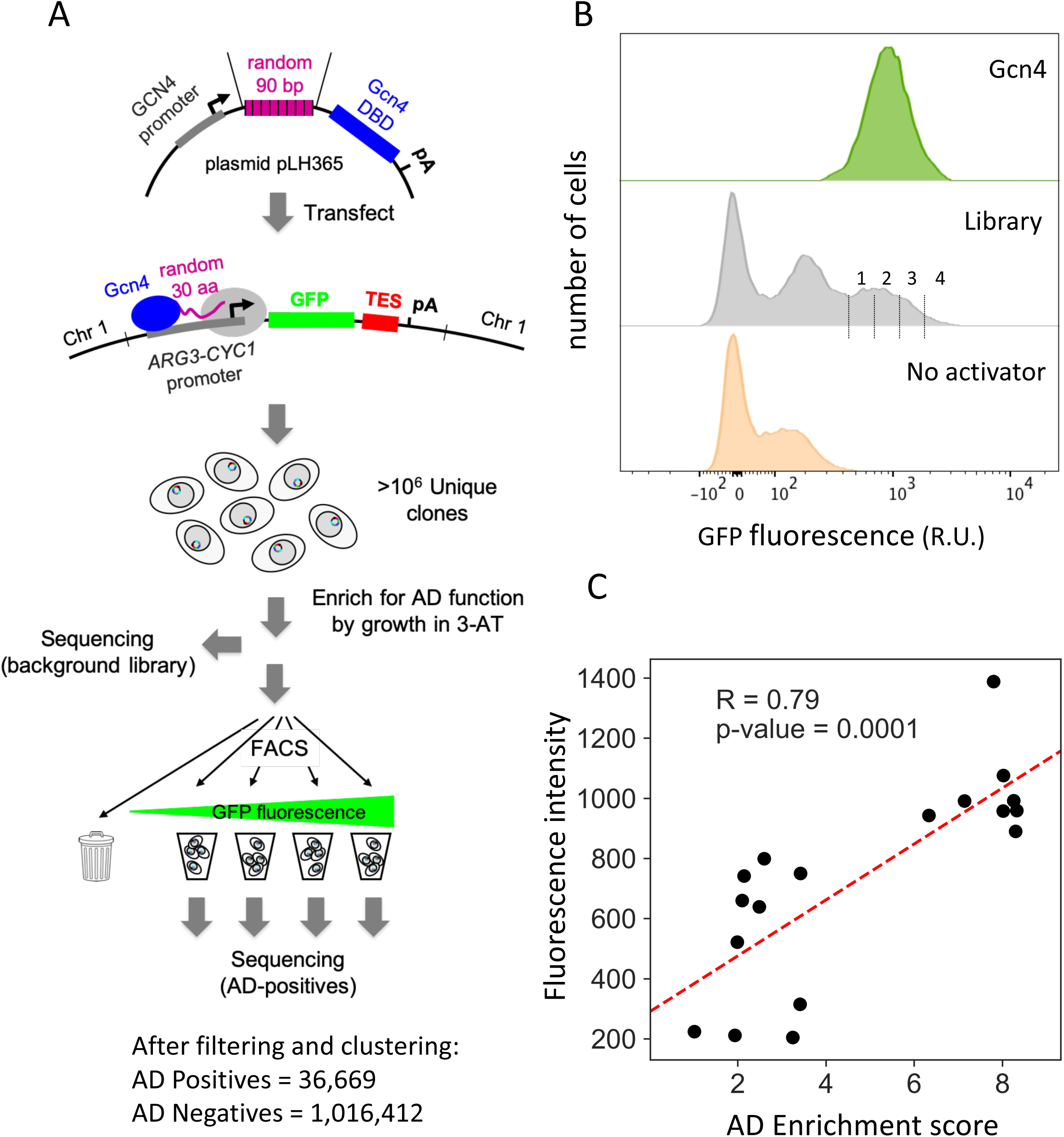
Experimental design and validation. **A)** Schematic of high throughput screen for ADs. Cells contain a library of random 90-mers fused to the N-terminus of the Gcn4 DNA binding domain and a GFP reporter driven by a synthetic Gcn4-dependent promoter. Cells with Gcn4-AD function were enriched by growth in 3-AT followed by FACS to isolate the GFP-positive AD-containing cells. DNA from the library before FACS and from the four GFP-containing bins were sequenced. TES: *ADH1* terminator; pA: poly-A site. **B)** Density plots of the number of cells vs relative fluorescence intensities from FACS analysis. The three plots show cells containing WT Gcn4, the 3-AT-enriched library and cells containing no Gcn4. Vertical lines in the middle plot show a schematic of the gates used for binning AD-containing cells. **C)** Experimental validation of enrichment scores on 18 AD sequences versus GFP expression in the reporter strain. Sequences were picked from a wide range of enrichment scores. The AD enrichment score measures enrichment of the AD sequence in bins 1 to 4 with respect to the background library.

The libraries were transformed into a yeast reporter strain lacking wild type (WT) Gcn4 and containing an integrated synthetic Gcn4-dependent promoter driving GFP expression. Approximately 25 million yeast transformants were obtained, and ∼3.6 million contained uninterrupted ORFs fused to Gcn4. To enrich for functional ADs, we grew cells overnight in synthetic media lacking histidine and containing 3-amino triazole (3-AT), a competitive inhibitor of the yeast His3 protein. *HIS3* transcription is stimulated by Gcn4, and only cells containing functional Gcn4 produce enough His3 to efficiently grow under these conditions (Hope and Struhl, 1986). After selection in 3-AT, we sorted cells by their GFP levels using fluorescence-activated cell sorting (FACS). The distribution of fluorescence intensities shows that a subpopulation of cells expressed GFP at levels near those of cells with wild type (WT) Gcn4 (**Fig 1B**). FACS was used to split these GFP-expressing cells into four bins of increasing fluorescence. We predicted that cells with the highest GFP levels (bin 4) should contain the strongest ADs. DNA was extracted from cells in the individual GFP-expressing bins, and the 90 nucleotides encoding the random peptides fused to the Gcn4-DBD were sequenced. Only sequences containing a complete 30-residue ORF were analyzed. Single point mutations and other sequencing-related artifacts were minimized by clustering similar sequences, allowing for up to 6 mismatches per sequence to be included in the same cluster (Methods). The most frequent sequence in the cluster was used as the cluster representative. The AD-negative set contains all random peptide sequences from the background library (before FACS screening) except for those peptide sequences identified in any of the bins 1-4. The AD-positive set consists of all sequences found in any of the bins 2-4. Sequences that were found only in bin 1 were omitted as we later found that these likely contained at least some false positives (Methods). As a result, we obtained ∼4×10^4^ unique AD-positive sequences and ∼1×10^6^ AD-negative sequences (**Table S1**).

Most functional ADs are not found in a single bin but are distributed among several bins, with the distribution presumably reflecting AD strength. To check the accuracy of our FACS-based screening, we first assigned an AD enrichment score to each AD-positive sequence. This score measures the weighted enrichment of a 30-mer sequence in bins 1 to 4 with respect to its number of occurrences in the library prior to FACS screening (Methods). Next, we selected 18 AD-positive sequences from a wide range of enrichment scores and measured GFP expression in the reporter strain by fluorescence assay. We found that the calculated AD enrichment score correlates well with GFP expression induced by individual AD candidates, validating our activator screen (**Fig 1C;** Pearson correlation R = 0.79).

### Amino acid composition and specific dipeptide sequences are important predictors of AD function

We first compared sequences from the AD-positive and negative sets by calculating a log-odds score for each sequence based on its amino acid composition. This score measures the similarity of the sequence’s amino acid composition to the composition in the AD-positive set versus the composition in the AD-negative set (Methods). We found that the AD-positive and negative sequences have distinct but overlapping amino acid compositions (**Fig 2A**). This finding is consistent with earlier results showing that IDPs and ADs are generally composed of low complexity sequences that are biased towards certain amino acids.

**Figure 2.**
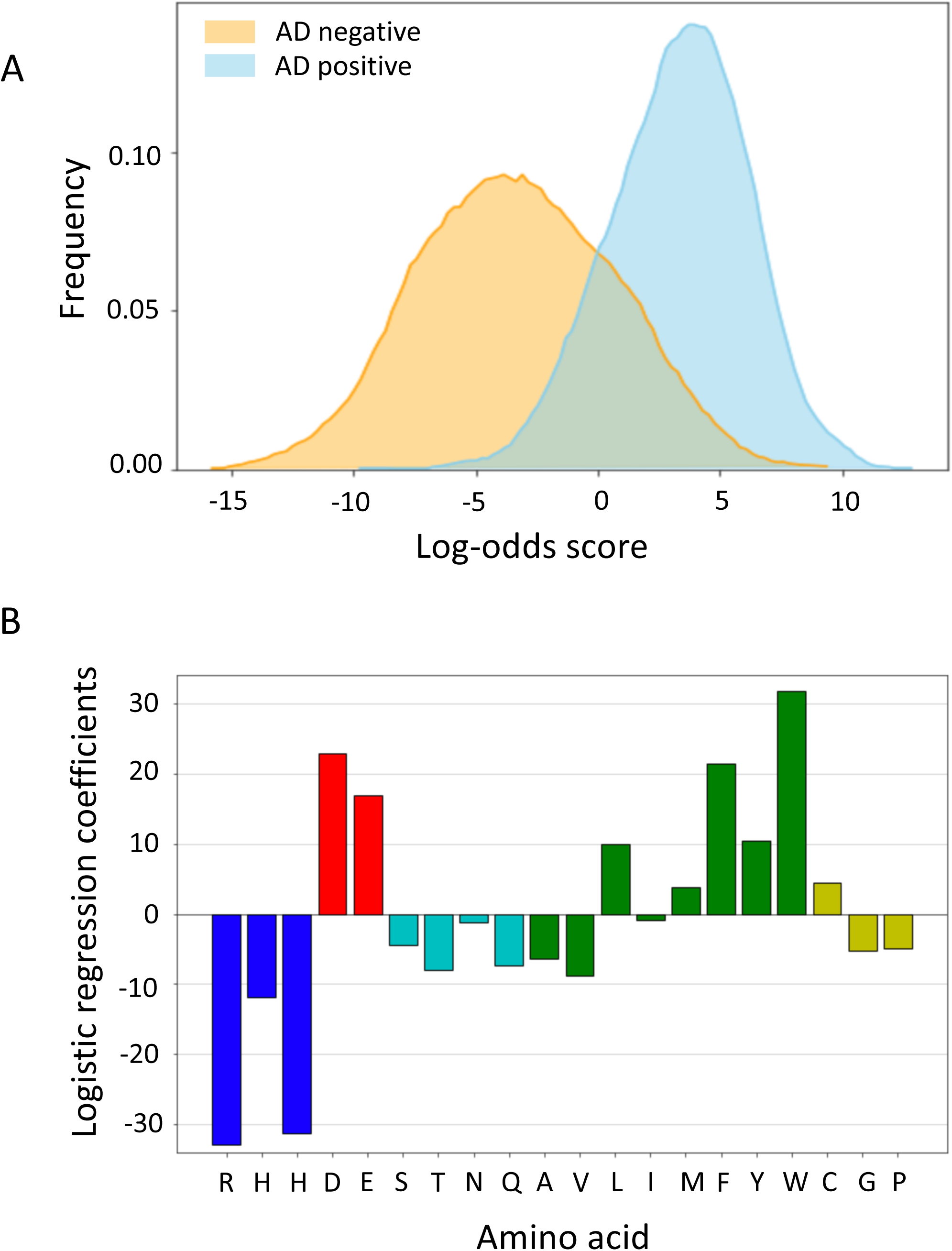
Properties of synthetic ADs. **A)** Distribution of log-odds scores for sequences from the AD-positive (blue) and AD-negative (orange) sets. **B)** Coefficients of amino acid frequencies derived from a logistic regression model for AD probability. Blue = positive charge; red = negative charge; green = hydrophobic/aromatic, cyan = polar and yellow = others.

To quantify the contribution of amino acid composition to AD function, we tested how well composition alone predicts function. We fit a logistic regression model for AD prediction that used only the relative amino acid frequencies (between 0 and 1) in each positive or negative sequence. The model was trained with 90% of the AD-positive and AD-negative data and tested with 10% held out data. Surprisingly, composition alone is a very strong predictor of function with an area under the precision-recall curve (AUPRC) score of 0.934 ± 0.002 (accuracy of predictions: 0.883 ± 0.003), compared with a maximum possible AUPRC of 1.0 for perfect predictions and 0.5 for random predictions. The logistic regression coefficients from this model show the bias towards specific residues in AD-positive sequences (**Fig 2B**). Consistent with results from prior analysis of natural and synthetic AD sequences (Cress and Triezenberg, 1991; Ma and Ptashne, 1987a; Pacheco et al., 2018; Ravarani et al., 2018; Warfield et al., 2014), the regression coefficients demonstrated that functional ADs are depleted of positively charged residues (R,H,K), and enriched for negatively charged (D,E), hydrophobic and aromatic residues, particularly F and W.

While no unique sequence or short linear motif has been recognized as conserved in natural ADs, it is possible that combinations of short heterogeneous sequence motifs contribute to AD function. To explore this possibility, we developed a regression model that utilizes the frequencies of all 400 possible dipeptide sequences. The resulting logistic regression coefficients from this analysis show the bias towards specific dipeptides that are enriched or depleted in the synthetic ADs (**Fig 3**). Using dipeptide instead of amino acid composition improved model performance from an AUPRC score of 0.934 ± 0.002 to 0.942 ± 0.002 (accuracy of prediction: 0.891± 0.004). Some dipeptides are clearly enriched in ADs such as D or E followed by a hydrophobic residue, especially F or W (log p-values from likelihood ratio tests are shown inside the boxes in Fig 3). The reverse dipeptides (e.g., W followed by D or E) show a negligible impact on the model performance (Methods). Importantly, we also found that certain dipeptides are strongly depleted in ADs, such as an aliphatic followed by a positive or polar residue, proline, or glycine (e.g. VP), whereas the same was not true for the reverse dipeptides (**Fig 3**). This analysis suggests that dipeptide sequences contribute to AD function over and above the contribution from their amino acid composition.

**Figure 3.**
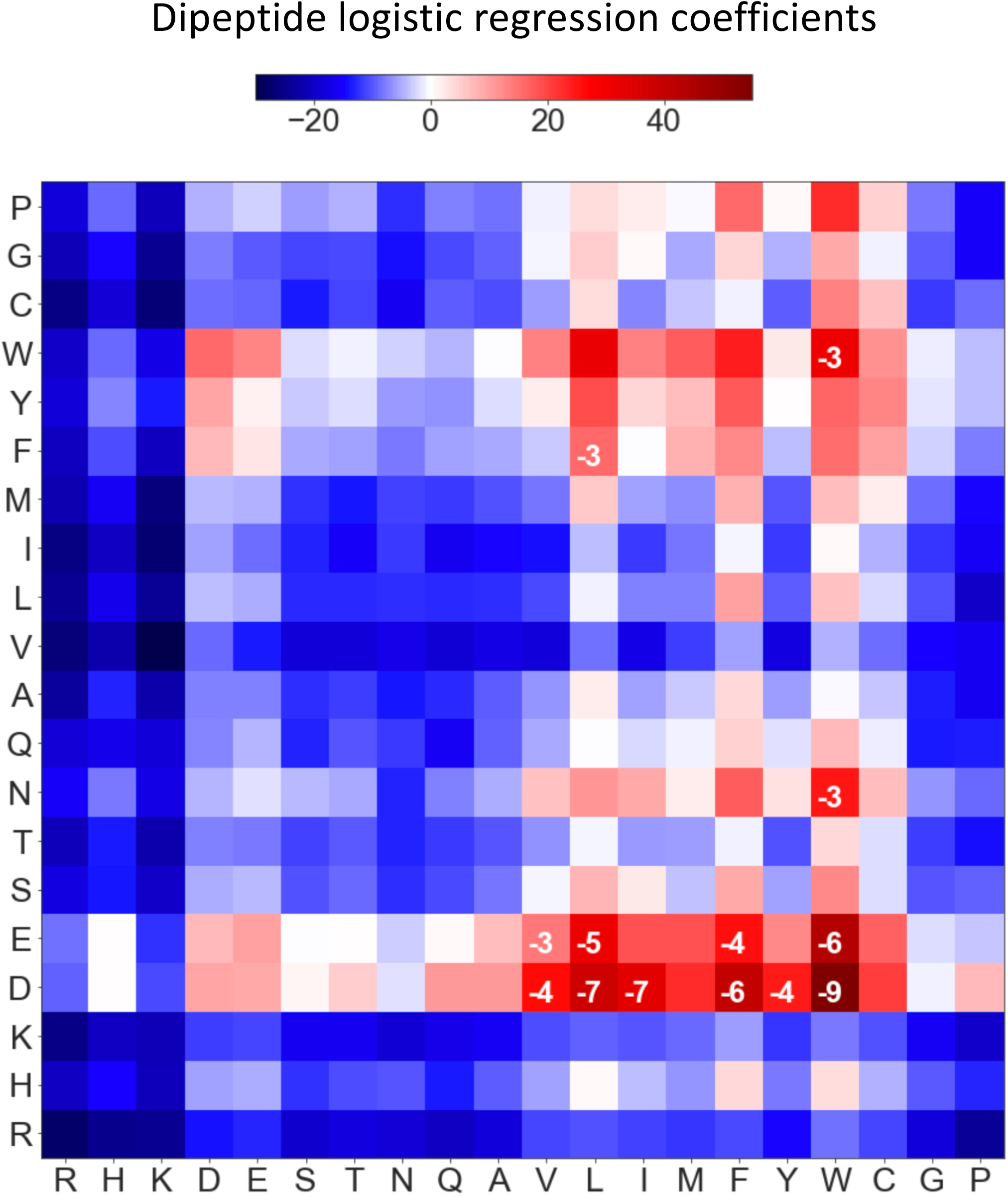
Dipeptide sequences contribute to AD function. Heatmap of the coefficients from a logistic regression model using only dipeptide frequencies. The first amino acid in the dipeptide is on the Y-axis. Log_10_ p-values are shown where p <0.001. p values are from likelihood ratio tests using all 400 dipeptide regression coefficients versus all but one (Methods).

To test these conclusions further, we swapped individual dipeptide coefficients in the regression model (e.g., the DW coefficient was swapped with all other coefficients in 400 separate models) and used the new models to predict AD function **(Fig S1A**). We found that replacing the DW coefficient (labeled Fwd) with every other coefficient in this matrix decreases average model performance significantly, while replacing the WD coefficient (labeled Rev) has no appreciable effect on average model performance. **Fig S1A** also shows that replacing coefficients for six similar dipeptides (EW, EV, DV, DL, DF, and DY) also decreased model performance while replacement of the reverse peptide coefficients does not. This analysis further indicates that these dipeptides contribute to AD function.

Since reversing dipeptides in the regression model affected our prediction performance, we asked whether the spacing between residues generally plays a role in determining activation function. We therefore compared occurrences of D-W separated by variable numbers of residues in functional and permuted sequences (**Fig S1B**). This analysis showed that at short spacings of 0-4 residues, having a D before a W is generally enriched over the opposite order. Altogether, our analysis showed that amino acid composition strongly correlates with AD function and suggests that enrichment of favorable dipeptide sequences and depletion of others contributes to function. These properties differentiate ADs from a control group of non-functional sequences.

We also compared the performance of our regression model with a previously proposed universal 9 amino acid AD sequence motif (Piskacek et al., 2007) (https://www.med.muni.cz/9aaTAD/) Both the regular and the more stringent 9aa sequence pattern did not perform well with our experimental data, achieving accuracies of 0.57 and 0.60, respectively.

### A deep learning model for AD prediction

To discover complex features that can contribute to AD function in an unbiased, agnostic fashion and to improve the accuracy of AD predictions, we trained a deep-learning neural network model that does not require prior knowledge of features contributing to AD function (Schmidhuber, 2015). The model inputs are the 30-residue sequences from each peptide in the positive and negative sets (20 values per position in one-hot encoding), predicted secondary structure (three values per position) and predicted disorder (one value) (**Fig 4A**). A series of 29 filters were used for data convolution that allowed us to model associations between residues at distant and variable positions. The resulting data is analyzed using a dense neural network with two soft-sign layers and the final output node yielding the probability of the input sequence to possess AD function. During training, the weights of the filters and other neural network connections are optimized, correcting for an imbalance of positives and negatives by subsampling the same number of negatives down to the same number of positives before each training epoch.

**Figure 4.**
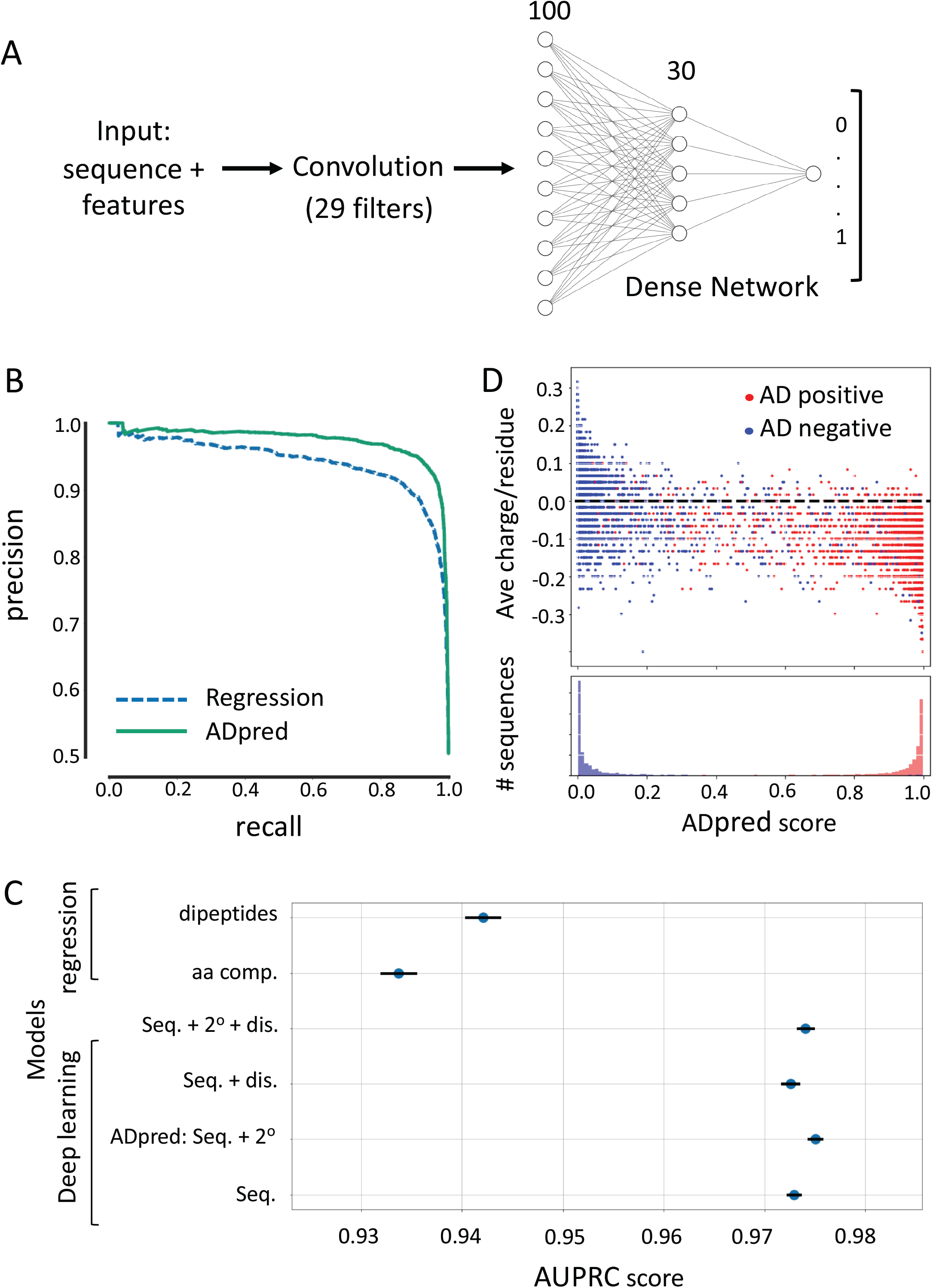
Deep neural network (NN) architecture and performance. **A)** Input to the deep NN is a 30 amino acid sequence and its predicted sequence features (secondary structure and/or intrinsic disorder). A convolutional layer can learn patterns characteristic of ADs independent of their precise position in the AD sequence. The flattened outcome of the convolution is used as an input for a dense two-layer-network with 100 and 30 neurons respectively. Lastly, the output layer composed of a single neuron gives the probability for the input sequence to have AD function. **B)** Analysis of model performance. The precision-recall curve compares the performance of the linear regression model utilizing dipeptide frequencies and the best deep learning model (ADpred) utilizing peptide amino acid sequences and secondary structure predictions. (**C**) Comparison of several regression and deep learning models evaluated with 10-fold cross validation, with the lines corresponding to standard error of the mean. Dis. = disorder predictions; Seq. = peptide sequence; 2^0^ = secondary structure prediction. **D)** Correlation between predictions of the deep learning model and the average charge per residue of the 30mers. Dotted line represents peptide with neutral average charge. Bottom panel shows a histogram for the distribution of sequences binned on ADpred probabilities.

**Fig 4B** compares the performance of the best deep learning and regression models. The best deep learning model, termed ADpred, uses only amino acid sequence and secondary structure predictions and shows great improvement in performance over the dipeptide regression model with an AUPRC score of 0.975 ± 0.001 (accuracy 0.932 ± 0.001) compared with an AUPRC score of 0.942 ± 0.002 (accuracy 0.891 ± 0.004) for the dipeptide model. We found that secondary structure but not disorder predictions modestly improved model performance (**Fig 4C**). The striking improvement in performance of the deep learning models over regression approaches suggests the existence of important features associated with AD function in addition to bias in amino acid composition and dipeptides sequences.

To evaluate the contribution of peptide charge for AD prediction using the deep learning model, we compared average charge per residue versus ADpred probabilities for both AD positive and negative sequences (**Fig 4D**). This analysis showed that extreme positive or negative charge correlates well with predictions, but many peptides cannot be accurately predicted by charge alone. For example, while we found few ADs with net positive charge, a large number of negatively charged peptides do not have AD function. This is consistent with our findings above that other features, in addition to amino acid composition, make important functional contributions.

### ADpred identifies functionally important AD residues

To test the utility of ADpred on a natural AD, we evaluated its performance on the Gcn4 central AD (cAD) where thousands of variants have been tested for *in vivo* function (Jackson et al., 1996; Staller et al., 2018; Warfield et al., 2014). We mutated a region of the cAD sequence *in silico* (residues 108-137) changing every residue to every other amino acid. We fed the resulting set of peptides to ADpred to predict AD probability. Remarkably, *in silico* predictions show excellent correspondence to prior results of *in vivo* function with a Pearson correlation of R=0.82 (**Fig. 5A, B**). For example, the importance of the three residues that make direct contact with Med15 (W120, L123 and F124; labeled in red in 5A) are clearly apparent as well as the lesser but noticeable impact of three other hydrophobic residues (F108, Y110 and L113; labeled in green) that have been observed *in vivo* (Jackson et al., 1996; Staller et al., 2018). Furthermore, our model predicts that insertion of positively charged residues are most likely to have a deleterious impact on function when positioned near the key hydrophobic residues, that insertions of additional hydrophobic residues generally increase function, and that no single negatively charged residue is important, in agreement with earlier *in vivo* studies (Jackson et al., 1996; Staller et al., 2018; Warfield et al., 2014).

**Figure 5.**
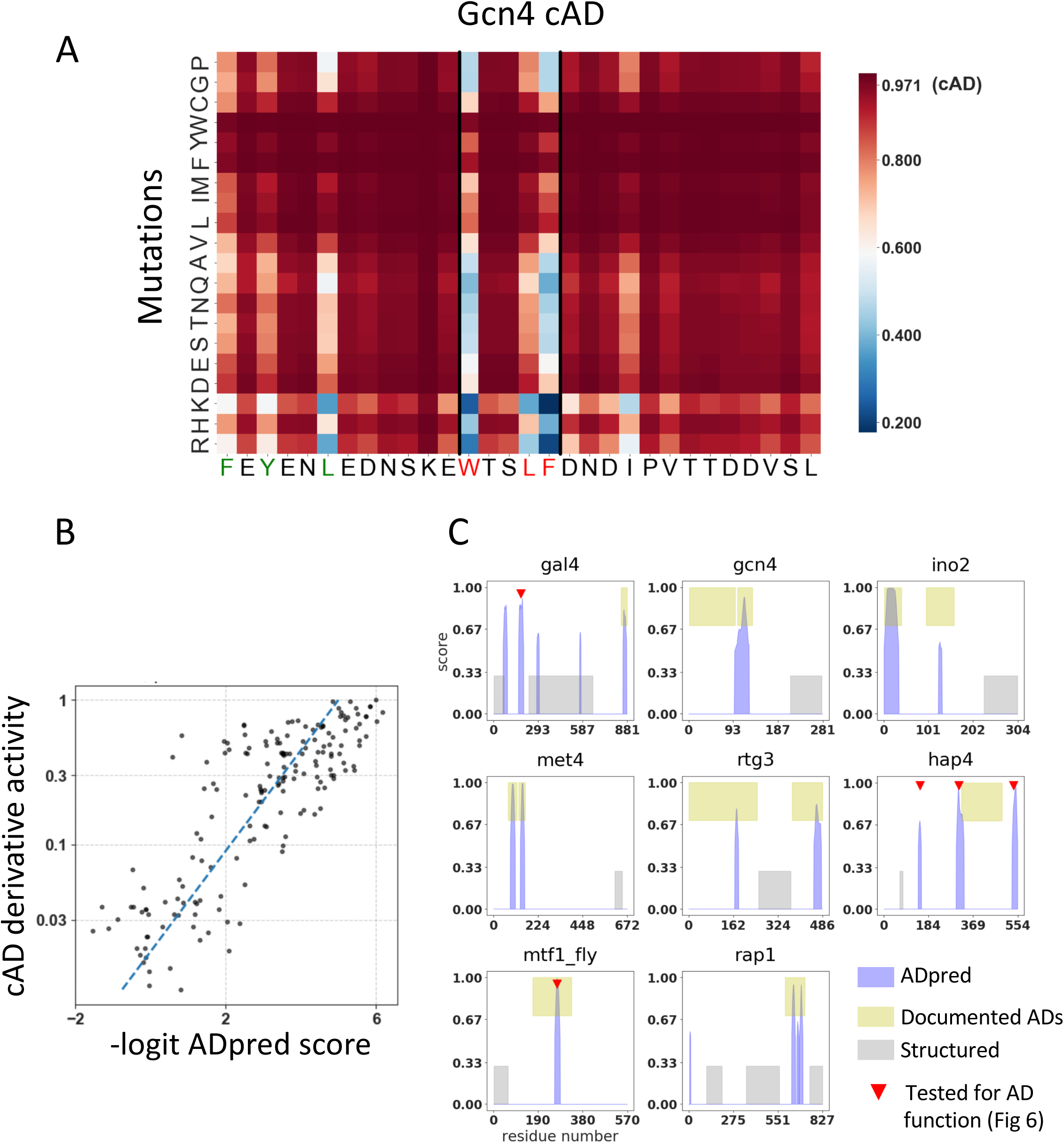
Performance of the deep learning model on known activators in yeast and fly. **A)** ADpred predictions of AD function probability for all possible single amino acid mutations of the Gcn4 central AD (cAD). An increase in ADpred score is darker red, decreases are lighter red or blue. **B)** The AD activity of the cAD constructs measured in (Warfield et al., 2014), transformed into 0-1 scale, shows a high correlation with the negative logit function of the ADpred probability (-log p/1-p). **C**) Several well-characterized activators from yeast and fly were analyzed by ADpred. Known activation domain-containing regions are colored green, ADpred predictions in blue and predicted structured domains in gray. Red triangles indicate predicted ADs tested for AD function (details in Fig. 6).

Using the same *in silico* mutagenesis approach, we predicted important residues within other natural ADs (**Fig S2**). Overall, we observe an excellent correspondence between the *in silico* predictions and the experimental identification of functional residues in the yeast activators Ino2 and Gal4 (Pacheco et al., 2018, Tuttle et al., 2019). However, two predictions deviating from the experimental results are Gal4 residues Y865 and Y867. Both are predicted to be functionally important, but when individually mutated to Ala they have at least 75% WT function (Tuttle et al., 2019).

We next used ADpred to identify ADs in transcription factors where *in vivo* AD function has been approximately mapped (**Fig 5C**) (Arnold et al., 2018; Kuras and Thomas, 1995; Leuther et al., 1993; Ma and Ptashne, 1987b; Pacheco et al., 2018; Rothermel et al., 1997; Schwank et al., 1995; Stebbins and Triezenberg, 2004). For this analysis, we used an ADpred probability of ≥ 0.8 as a high confidence threshold for AD prediction. In the figure, blue peaks show the AD predictions, while green boxes indicate experimentally validated AD function. For many factors (Gal4, Gcn4, Ino2, Met4, Rtg3, Rap1), our model predicts AD function coincident with or directly adjacent to known ADs. One exception is the Gcn4 N-terminal AD, where optimal AD function requires a combination of four short hydrophobic clusters scattered throughout the 100 aa long N-terminal region (Jackson et al., 1996; Tuttle et al., 2018). None of these four short clusters can act as an AD on its own but requires the others for its collaborative function. It is likely that our model could not find this long AD region because it was trained on peptides of 30 residues.

For some of the factors analyzed (Gal4, Hap4, Rap1), our predictions fell outside of the AD regions mapped *in vivo* and sometimes within segments having predicted structure (grey boxes) e.g., Gal4. One possibility is that these AD predictions are false positives, meaning that these peptides do not encode AD function. An alternative explanation is that these peptides have the potential for AD function but are not positioned in the proper context to function in their natural setting; e.g. are in structured regions. To investigate these scenarios, we tested several of these predicted ADs. 30-residue segments containing predicted yeast ADs (indicated by red triangles in Fig 5C) were fused to the Gcn4 DNA binding domain. Function was assayed *in vivo* by treating cells with sulfometuron methyl (SM) for 90 min to simulate amino acid starvation and to induce synthesis of Gcn4, followed by RNA quantitation using RT qPCR (**Fig 6A; Fig S3, Table S2**). When assayed at the Gcn4-dependent *HIS4* gene, a predicted AD from Gal4 (Gal4_A) (Ma and Ptashne, 1987b) and three from Hap4 (Hap4_A,B,C), produced 3.7- to 8.6-fold higher transcription compared with SM-treated cells lacking Gcn4 (labeled “vector” in Fig 6A; dashed line). At *HIS4*, we used activation of transcription by ≥ 3-fold for scoring AD+ function. Our results show that these predicted ADs do not inherently lack activity but can function as ADs in an appropriate context.

**Figure 6.**
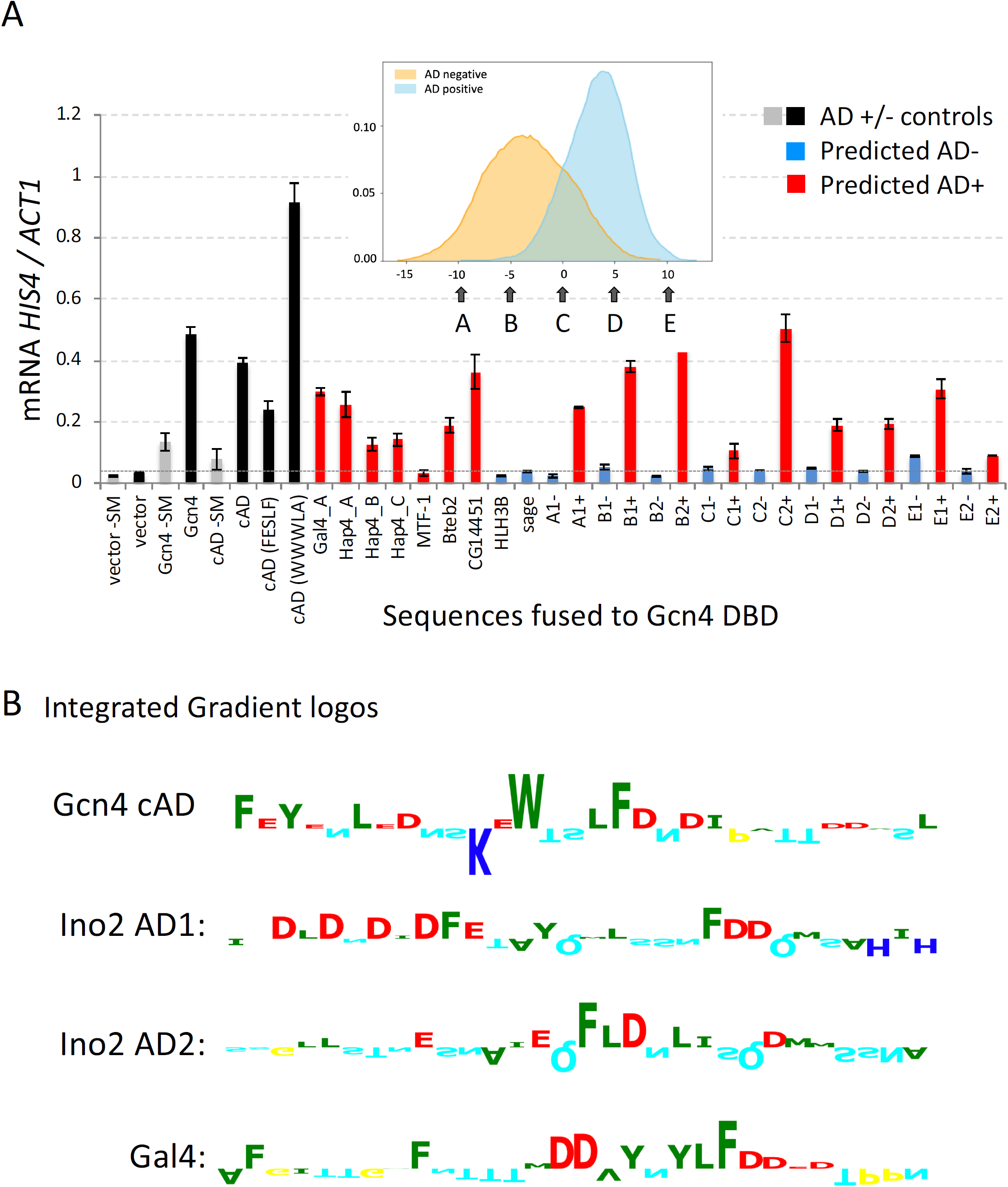
Performance of the deep learning model on predicted natural and synthetic ADs. **A**) RT qPCR quantitation of mRNA from the yeast *HIS4* gene, normalized to *ACT1* mRNA. Cells contained the indicated AD sequence (**Table S2**) fused to the Gcn4 DBD in pLH365 and were induced with SM for 90 min before mRNA quantitation. Grey bars = no SM added, all others have SM; Black bars = control Gcn4 derivatives including WT Gcn4, the Gcn4 cAD and two previously characterized cAD derivatives (FLESLF and m115) (Warfield et al., 2014). Red bars = sequences predicted to have high probability of AD function; blue bars = low probability of AD function (Fig S3B). Dotted horizontal line indicates the level of SM-induced transcription in cells lacking Gcn4. **Inset:** the histogram of log-odds scores is from Fig 2. Arrows point to regions where selected AD-positive sequences were randomized and used to search for one or two pairs of sequences with the same amino acid composition and high (+) or low (-) ADpred scores (Methods). Labeling of sequence pairs with identical log-odds scores is with a letter corresponding to Fig 6A inset followed by 1 or 2; e.g., A1+/A1-, A2+/A2-, etc. **B)** Predicted importance of individual residues for ADpred scores analyzed using Integrated Gradients. Contributions (positive upwards, negative downwards) of the residues in 4 selected yeast ADs (colors are the same as in Fig 2A).

To test the performance of our model on activators from other eukaryotes, we measured the AD function of several sequences analyzed in *Drosophila* using a medium-throughput AD-screening approach (Arnold et al., 2018). We examined sequences from factors MTF-1, Bteb2, and C14451 that were shown to have AD function in *Drosophila* and were predicted to have AD function using ADpred (**Fig S4. Table S2**). While MTF-1 did not function as an AD in our system, the other two had AD activity resulting in 5-10-fold higher levels of transcription compared with cells lacking Gcn4 (**Fig 6A** and **Fig S3**). We also tested regions of two factors, HLH3B and Sage, that have AD function in *Drosophila* but were predicted by ADpred to have no AD function. We found that neither of these sequences functioned as ADs in the yeast system, further validating the ADpred results.

### ADpred can overrule strong amino acid composition bias

As demonstrated above, amino acid sequence composition is perhaps the most important component determining AD function, but other features also make important contributions. Given that a model using only sequence composition as a feature reaches quite high accuracies, we asked whether the ADpred predictions are dominated by sequence composition. We selected sequences from our libraries within a wide range of log-odds scores for amino acid composition (labeled A to E in **Fig 6A**, inset). For each selected sequence, we generated a set of 10,000 randomly permuted 30-mer peptides and then sorted them using ADpred. From this set, we selected one or two pairs of sequences with identical amino acid composition but with high and low ADpred scores (AD+ or AD-). Upon testing these pairs of 30-mers for function at *HIS4* and using activation of transcription by ≥ 3-fold for scoring AD+ function, all predictions were confirmed except for one of two sequences tested with +10 log-odds score (**Fig 6A**). Sequence E2+ has a sequence composition extremely biased toward AD function but only shows 2.6-fold activation (**Fig 6A**). Combined, our results demonstrate that ADpred can correctly predict AD function with high accuracy even if the sequence composition is strongly biased toward non-AD sequences and vice versa.

We also tested all of the above peptides for activation of yeast *ARG3* transcription (**Fig S3A**). *ARG3* is transcribed at ∼7-fold lower rate compared to *HIS4*, and transcription of *ARG3* is regulated by both Gcn4 and two repressors. Our prior studies using AD derivatives at both promoters showed that *HIS4* is generally more permissive for AD function, perhaps because of the more complex regulation and coactivator requirements at *ARG3* (Pacheco et al., 2018; Tuttle et al., 2018). Because WT Gcn4 shows lower levels of activation at *ARG3* compared with *HIS4* (5.5-fold vs 14-fold), we set a threshold of 2-fold activation for scoring AD function (**Fig S4**, dashed line indicates no Gcn4). Of the four AD predictions for yeast proteins outside of previously mapped ADs, only one activated *ARG3* >2-fold (Hap4A), but four of five *Drosophila* proteins and 15 of 18 synthetic sequences behaved as expected. Thus, our predictor performs well but is less accurate on a promoter with more stringent AD requirements (74% accuracy at *ARG3* compared to 93% at *HIS4*. Nevertheless, there is a high correlation of experimental vs. predicted values at both *HIS4* and *ARG3* with R=0.85 and 0.67 respectively (**Fig S3B**).

### ADs generally contain clusters of hydrophobic residues rather than specific sequence motifs

As demonstrated above, ADpred performs significantly better than regression models based on amino acid composition and dipeptide frequency. For additional insight into sequence features leading to enhanced performance of the neural network, we analyzed the results using Integrated Gradients, a method that identifies positive and negative contributions to the prediction score (Ancona et al., 2018; Sundararajan et al., 2017). This analysis of four representative yeast ADs is shown in **Fig 6B**, with the results presented as sequence logos. In addition, Integrated Gradient analysis of 20 high-scoring synthetic peptides is shown in **Fig S5**. In contrast to earlier predictions (Piskacek et al., 2007; Warfield et al., 2014), we found no evidence for ADs to contain defined sequence motifs of three or more residues. Rather, a common feature is clusters of hydrophobic residues in the background of an acidic polypeptide. This result explains the importance of amino acid composition, as multiple examples of this simple sequence feature are likely to be found within short peptides having a favorable amino acid composition.

We also used Integrated Gradients to examine the scrambled peptides with variable amino acid composition used in Fig 6 (**Fig S6**). This analysis showed that peptides with predicted ADs, but having unfavorable amino acid composition, had separately clustered the favorable and unfavorable residues. For example, peptides with compositions labeled A and B had positive residues segregated to the N-terminus while the acidic and aromatic residues were positioned in the C-terminus. This further validates our conclusion that clusters of hydrophobic residues in the background of an acidic polypeptide are important for function.

### *ADs show higher helical propensity and less disorder than surrounding sequences* from *in silico* analysis

To further explore properties of natural ADs, we applied the deep learning model to the entire yeast proteome. We classified protein regions as AD+ or AD-if a segment of five or more residues above a high-confidence threshold ADpred probability of 0.8 was detected. We compared predictions within the proteome to predictions on a subset of 132 yeast transcription factors, some of which are known activators (**Fig 7A, Table S3**). We observed a clear enrichment of ADs in transcription factors, and a corresponding low *p-value*, when the predicted AD is between 15-30 residues in length. While predicted ADs are enriched in transcription factors, they are still found, but at a lower level, in non-transcription factors. This is in agreement with our findings above that not all peptides with inherent AD function are in a context that allows them to function as activators.

**Figure 7.**
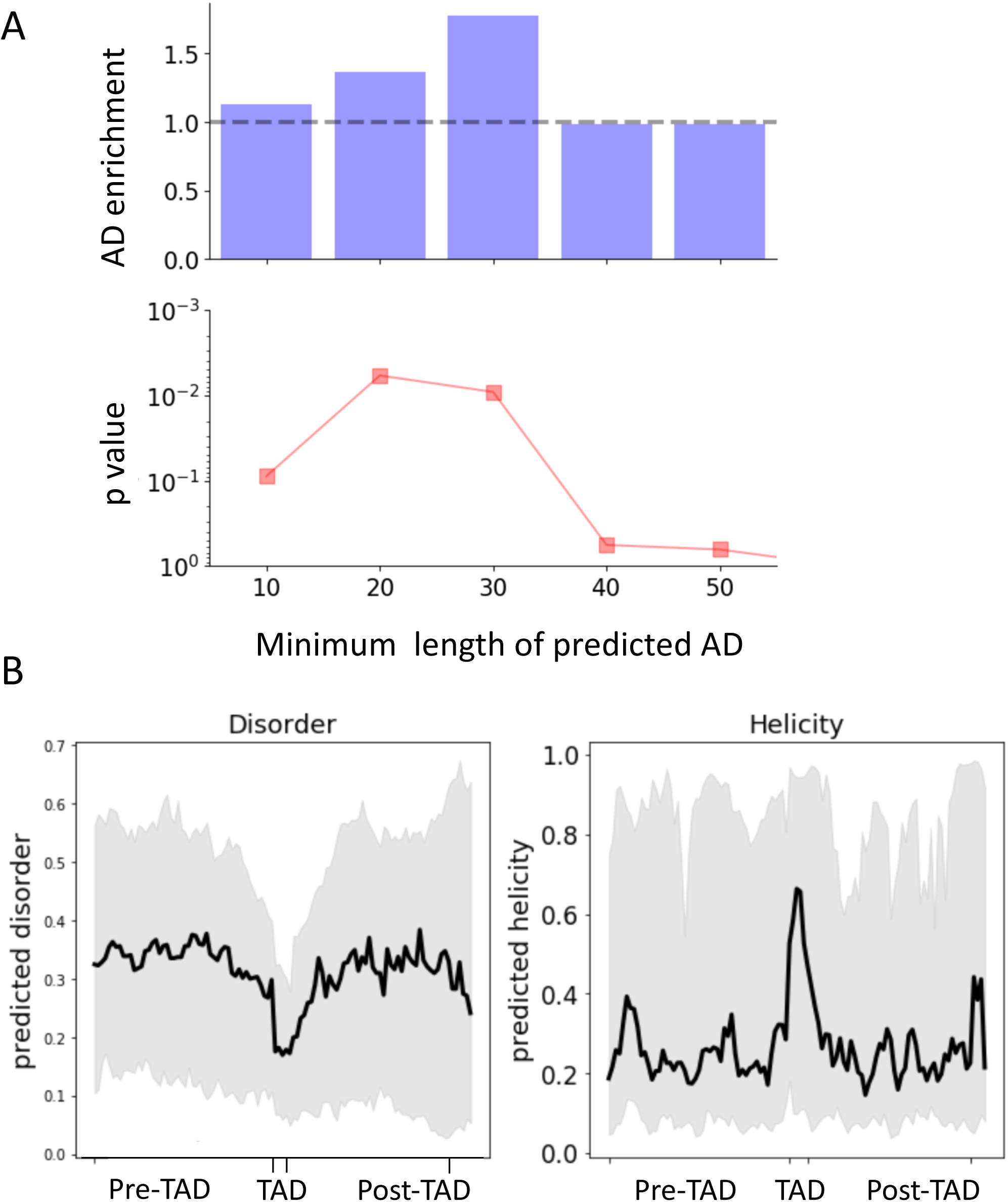
Properties of predicted AD regions in yeast transcription factors. **A**) Enrichment of predicted ADs in yeast transcription factors (**Table S3**) compared to the complete proteome plotted against the minimum length threshold for calling an AD. The enrichment p-values from a Fisher test are shown below. **B**) Predicted disorder and helicity in and around predicted ADs in a set of 132 yeast transcription factors (**Table S3**). Black thick line: median values; Grey: values between the 25th and 75th percentile.

Finally, we explored whether sequences within the proteome having predicted AD function are enriched for disorder or secondary structure elements. Our analysis examined the 25, 50 and 75th percentiles of the predicted helical propensity or disorder within 50 residues N and C-terminal to the predicted AD and five central residues of the ADs (the latter is to plot ADs vertically stacked, despite their variable length). The predicted ADs have, on average, lower disorder compared to the surrounding sequence (**Fig 7B**). Consistent with this, ADs are also predicted to have higher alpha helical propensity compared with 50 residues N and C-terminal to the predicted AD. Together, our analysis suggests that many natural ADs are peptides with alpha helical propensity located within disordered regions. We found this same pattern whether analyzing the entire yeast proteome, the subset of nuclear proteins, or only yeast transcription factors (**Fig S5**). We therefore suggest that the observed pattern of helicity and disorder might be some inherent property of the “AD-type peptides” and their normal protein environment, whether or not they are transcription factors.

## Discussion

Since their discovery, the nature of transcription activation domains has been enigmatic (Gill and Ptashne, 1987; Hope et al., 1988; Ma and Ptashne, 1987a; Ptashne and Gann, 1990; Sigler, 1988). Nearly all characterized ADs are intrinsically disordered and have biased amino acid composition. Furthermore, function corelates with negative charge and in some cases, with the length of the AD region. Earlier studies showed that a surprisingly large percentage of random sequences can encode AD function, and examination of natural and variant ADs revealed no obvious common sequence motif. Despite these unusual properties, activators typically target only a small number of conserved coactivator targets, and the individual AD-target interactions usually have relatively low affinity and specificity. Recent structural studies showed that one large class of ADs interacts with its targets via a dynamic fuzzy interface and that combining multiple ADs and activator binding domains in a single complex can lead to much higher affinity and specificity while still retaining the dynamic and fuzzy nature of the complex. However, major hurdles toward understanding AD function and specificity are the absence of a reliable way to recognize ADs in transcription factors and an understanding of the protein features that permit their distinct functions. Here, we used a high throughput assay to screen over a million synthetic peptide sequences for their ability to activate transcription of a GFP reporter. We analyzed the resulting sequence sets using logistic regression and a deep neural network that can accurately predict AD function. The combination of these two approaches allowed us to identify, in a systematic way, properties and sequence features associated with AD function.

The starting point for our analysis was a high throughput screen for ADs that included a genetic enrichment for ADs within the randomized library. Although randomized libraries have been screened for AD function in earlier work (Abedi et al., 2001; Erkine et al., 2002; Ma and Ptashne, 1987a; Ravarani et al., 2018), compared with prior screens, we identified ∼60-fold higher numbers of ADs and a much larger number of non-ADs, an important starting point for systematic analysis of functional properties. For example, an earlier machine learning approach used 926 synthetic AD variants that gave an AD prediction AUROC score of 0.773 (in comparison to our AUROC of 0.977; **Table S4** in Supplementary Methods) and, significantly, attributed different relative importance to some of the features described here (Ravarani et al., 2018).

As inferred from earlier studies, we found a striking difference in amino acid composition between the AD-containing and non-AD sequences. A logistic regression approach based solely on amino acid composition was surprisingly accurate (AUPRC 0.934). Regression allowed us to quantify the contribution of residue type to predicted function, and this was consistent with earlier work: ADs are generally depleted of positively charged residues and enriched for acidic, hydrophobic and especially aromatic residues. This approach also allowed us to examine the contributions of simple sequence motifs. Our analysis showed that ADs are enriched for specific dipeptides and depleted of others. One of these dipeptides, DW, had been identified earlier (Ravarani et al., 2018). This is in agreement with a prior proposal that one function of acidic residues in ADs is to promote solvent exposure of hydrophobic residues that are involved in direct molecular interactions (Staller et al., 2018).

To further improve performance and to enable predictions on a proteome-wide scale, we developed a deep neural network for AD prediction. Deep learning allows predictions of function without *a priori* knowledge about which patterns or properties might be important for the prediction. These properties include short and variable sequence motifs and residue spacing, both internal and between multiple short motifs that positively or negatively contribute to function. Using our large data set of AD positive and negative sequences, with the addition of only secondary structure predictions, this approach gave a striking improvement in the accuracy of AD prediction compared to the logistic regression model (AUPRC 0.977 compared to 0.934). We also found that the deep learning model, termed ADpred, performs well, even with sequences that show extreme bias in amino acid composition against AD function. Including features representing predicted disorder did not increase performance. This is not surprising, since it is unlikely that a sequence in our random library would by chance fold by itself.

Analysis of sequence features that contribute to the ADpred results using Integrated Gradients showed that many of the ADs contain clusters of hydrophobic residues in the background of an acidic polypeptide. This feature is found in both natural and synthetic ADs and seems a general feature corresponding to function. We suggest that these clusters function to increase the effective affinity of the AD peptides for their coactivator targets using a mechanism similar to avidity or allovalency – whereby a receptor dynamically interacts with multiple binding sites on a single ligand, effectively inhibiting the dissociation of the two molecules (Locasale, 2008; Olsen et al., 2017). This mechanism fits nicely with the dynamic and fuzzy binding of acidic activators to Med15, and presumably other coactivator targets, as well as the finding that AD-coactivator binding is driven in part by a favorable entropy change (Pacheco et al., 2018; Tuttle et al., 2018).

Our new results, combined with earlier work, suggest that functional ADs (1) consist of a disordered polypeptide with biased amino acid composition, (2) contain clusters of hydrophobic residues in the background of an acidic polypeptide, (3) are enriched for specific short dipeptide sequences and depleted of others, and (4) have less disorder and more helical propensity than surrounding sequences that facilitate the presentation of their hydrophobic residues to interacting partners. Taken together, our characterization fits with a fuzzy-binding mechanism where the interactions take place in a dynamic environment resembling a hydrophobic cloud rather than combinations of sequence-specific interactions.

Tests of our optimized model showed that it can accurately identify functionally important residues within yeast ADs. For example, *in silico* mutagenesis of the Gcn4 cAD to every possible residue and predicting the effect on AD probability gave results remarkably consistent with the extensive experimental analysis of Gcn4 AD function. When applied to several factors with known ADs, the model was successful in AD recognition, with the exception of the highly cooperative 100-residue long Gcn4 N-terminal AD – a polypeptide length far outside the parameters of our model training. We also found potential ADs within other regions of transcription factors. In most cases tested, these predicted ADs functioned when fused to the Gcn4 DNA binding domain. This result demonstrates that AD function requires that the polypeptide be located in the proper protein context and that not all sequences within natural proteins having AD potential will work as activators. Recognition of these “false-positives” when screening the proteome will require additional information. For example, ADpred was trained on short random sequences, which are likely to be disordered. Identification of true ADs in transcription factors will likely be more accurate if only disordered regions are considered (**Fig 5C**).

Finally, it is important to note that our screen used a TATA-containing inducible promoter. Earlier studies have shown that enhancers, the DNA targets of activators, can have specificity for a certain promoter type and that coactivator requirements can vary dependent on the gene regulatory region (Butler and Kadonaga, 2001; Haberle and Stark, 2018). Differences in coactivator requirements probably play a role in the differences in function when comparing synthetic ADs on *HIS4*, a more permissive gene, and *ARG3*, a more difficult gene to activate. This may also account for the differences we observed for the function of some *Drosophila* ADs in the fly and yeast systems.

Some yeast acidic activators, such as Gal4, work in all eukaryotes, and the ADs we have isolated here have similar properties and are likely of this class. In contrast, some higher eukaryotic cell-type specific activators bind particular coactivator targets using a sequence-specific and conventional protein-protein interface that likely have different sequence requirements, e.g., (De Guzman et al., 2004). It will be of great interest in future work to repeat the screen using promoters with different coactivator requirements and promoter sequence elements to determine whether this setup changes the features necessary for AD function. In addition, it will also be of interest to test how predictions of AD function correlate with the ability to form condensates – a property associated with at least some ADs (Hahn, 2018).

## Methods

### Design of the randomized libraries

For the first library, we computed the ratio of A, C, G, and T needed at each codon position in order to obtain an approximately equal probability for encoding each of the 20 amino acids and a minimal probability for a stop codon within our random 90-mer nucleotide sequence. In a custom python script we minimized an objective function using the “basin hopping” algorithm (Wales and A, 1997) implemented in the Python scientific library scipy (Oliphant, 2007). The objective function is the Euclidean distance from equal representation of all 20 amino acids (Pr(aa)=0.05) and an absence of stop codons (Pr(stop)=0). The goal for the second library was to obtain amino acid target probabilities equal to the average observed in disordered regions. We used the same Python script to compute the optimal ratios of A, C, G, and T to minimize the Euclidean distance with target and predicted probabilities.

Oligonucleotides containing 30 repeats of the randomized codons (Supplementary Methods) were ordered from Integrated DNA Technologies (Coralville, IA). Each of the three codon positions contains a defined ratio of A/C/G/T to generate the desired bias. The oligonucleotides were extended in a PCR reaction to add 40 bp identity on each end to plasmid pLH365 (Supplementary Methods). This plasmid was derived from the ARS CEN *LEU2* vector pRS315 and contains 1 Kb upstream DNA and the coding sequence for Gcn4 residues 132-281. This upstream DNA contains all known Gcn4 promoter and translational regulatory elements. The plasmid was digested with SbfI and NotI and 4 μg of linearized vector, and 12 μg of the PCR products were transformed to electrocompetent yeast strain SHY1018 so that in vivo homologous recombination inserted the randomized 30-mers into the N-terminus of Gcn4 (Benatuil et al., 2010). Ten transformations were run in parallel to produce a library of ∼2×10^7^ clones. The data from both libraries were combined into a single dataset to increase the number of samples and improve model performance.

### GFP induction and FACS sorting

Cells were diluted to OD_600_= 0.3 from a suspension with OD_600_= 1.0 and grown overnight at 30°C in glucose complete media lacking uracil, leucine and histidine and containing 3 mM 3-Amino Triazole (3AT). After 14-19 hours, cells were washed and diluted in double distilled water to ∼10^7^/ml and FACS sorted in a FacsAriaII instrument. Cells were sorted into 4 bins of increasing fluorescence intensity and collected. Sorted cells were harvested by centrifugation and resuspended in glucose complete media lacking leucine and containing 100 μg/ml ampicillin and incubated for 2 days at 30°C with shaking. Cells were frozen in 20% glycerol at −80°C.

### Plasmid purification and High-throughput sequencing

The background libraries were obtained from cells after growth in selection media but before FAC sorting. Cells in FACS-sorted bins 1-4 were barcoded using Illumina nextera i7 barcodes and sequenced on a Illumina HiSeq machine. For each library, approximately 10^9^ cells from a suspension with OD_600_ =1.5 were harvested by centrifugation, washed with water and resuspended in 4 ml of TE buffer. Cells were lysed in a mini bead beater (BioSpec products) with ml of zirconium beads 7 times at maximum speed for 3 minutes and with rest intervals of 5 minutes on ice. DNA was extracted with Phenol/CHCl_3_, ethanol precipitated and treated with RNAseA. The libraries for high-throughput sequencing were prepared as described in Supplementary Methods.

### Data processing for machine learning

All procedures are implemented in custom python and bash scripts. Reads 1 and 2 from paired-end sequencing were paired with FLASH (Magoc and Salzberg, 2011). We filtered out sequences longer than 90 base pairs, with sequencing quality PHRED score less than 30 for a given base, with frameshifts, or without start or stop codons. Paired nucleotide sequences were translated to amino acids. Sequence clustering (Edgar, 2010) was applied to minimize redundancy in the libraries (with minimum sequence identity of 80% per cluster). Each cluster was represented by its most frequent member sequence. Each such sequence is included in the final reduced dataset and the total number of reads in bins 1 to 4 and background correspond to the sum of reads of all members of the cluster. For an initial experimental validation, an activator enrichment score was calculated for each sequence in the final dataset as the summation of the number of reads in each bin, multiplied by coefficients (coeff_bin_) that correspond to the mean value of fluorescence of each bin (Supplementary Materials):

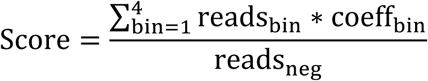

Here, reads_neg_ stands for the number of reads in the AD-negative set, which comprises all reads from the unsorted background library minus any AD sequences found in FACS bins 1 to 4.

### Machine learning analysis

The redundancy-filtered set of sequences with read counts in bins 1 to 4 and background were split into positive (AD-positive) and negative (AD-negative) sets. The AD-negative set contains all sequences in the background library except those found within FACS bins 1 to 4. The positive set contains all sequences with at least one read in total in bins 2 to 4. Omitting sequences that were only found in bin 1 improved model performance, presumably by eliminating false positives. Charges were computed for each sequence as the summation of amino acid frequencies multiplied by a coefficient where (E,D=-1; K,R=1 and H=0.5).

For **Fig. 2A**, we compared the sequence composition between the positive and negative sets by computing the log-odds score for each sequence and plotting its distribution for the two libraries. The log odds score for each sequence was calculated as the sum of the log enrichments of each of the 20 amino acids in the sequences, where enrichment is the ratio of amino acid frequencies in the AD-positive versus the AD-negative sequence sets. Positive scores indicate an amino acid composition similar to the AD-positive set while negative scores indicate a composition similar to the AD-negative set. Denoting with 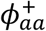 the averaged frequency of the amino acid aa in the positive set and with 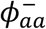 the frequency in the negative set, the log odds score is:

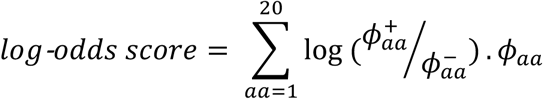

In Fig. 2B, we trained a logistic regression model with L2 regularization (*λ* = 3.9 10^−3^ was chosen from a grid of 40 default values provided in *LogisticRegressionCV* function from scikit-learn package) to predict whether a sequence is AD-positive or AD-negative. The model was evaluated using 5-fold cross-validation. This required optimizing 21 parameters, one for each amino acid frequency in the sequence and one offset. For Fig. 3, we trained a logistic regression model to predict AD function using the 400-dimensional dipeptides composition instead of the 20-dimensional single amino acid composition.

To assess the importance of specific dipeptides for AD function, we performed 400 likelihood ratio tests, each comparing the full model with models lacking one dipeptide feature. Dipeptides with significance p-values below 0.001 are indicated in Fig. 3 by their base 10 logarithm. We also built alternative AD score distributions by flipping the coefficients of a dipeptide to every other dipeptide and measuring the performance of these models in the test and validation datasets (**Fig S1A**). We repeated this for all dipeptides, or the specific dipeptides shown, and compared the distribution visually with boxplots.

The deep neural network for ADpred (**Fig 4A**) was implemented using Keras 2.1.6 (Chollet, 2015) with a TensorFlow (tensorflow.org) backend. Briefly, the input was composed of sequence and secondary structure (H, E and –, for Helix, β-sheet and random coil from PSIPRED 4.0.1) in a one-hot encoded matrix of dimension 30 by 23. This input was fed into a model made up of a convolutional layer, two dense layers and the output dense layer. The convolution layer had 29 filters with filter size 6×23, the first hidden dense layer had 100 neurons and the second hidden layer had 30 neurons. Each layer had a softplus activity (log(1 + *e*^*x*^)). The hidden layers were regularized with L2 regularization (λ=0.001) and dropout (P=0.5) layers. The final output layer had a single neuron with a sigmoid activation function and was used to compute the final probability for AD function prediction.

The model was trained with the ADAM optimizer (Kingma & Ba, 2014), using the binary cross entropy loss function, and the model’s performance was analyzed using AUPRC (area under a precision-recall curve), which corrects for skewed class sizes and is a common metric used in classification tasks. Each epoch was split into 250 batches. At the beginning of each epoch, we randomly drew an equal number of positive and negative samples from the original data set.

The performance of ADpred was evaluated using 10-fold cross-validation. This process involved using 8 of the folds for training, 1 of the folds for validation, and the final fold for testing. The fold used for testing rotated between the ten folds over ten iterations. At each iteration the model hyperparameters (batch size, number of epochs, optimization algorithm, learning rate and momentum, activation functions, drop out probabilities and convolutional filter size) were determined using grid search over a pre-defined set of possibilities. To reduce variance, 10 random weight initializations were used for each set of hyperparameters. The model that performed the best on the validation set out of all combinations of hyperparameters and weight initializations was subsequently evaluated on the test set. The evaluation metric on both the validation set and the test set was average precision, which is a conservative approach for calculating the area under the precision-recall curve. The results are shown in Figure 4B To train ADPred, each hyperparameter was fixed to the mean of the optimum over its 10 values (detailed in the previous paragraph). Then the complete set was split again into 1 part as a test set and 9 parts for training. The best model over 100 random initialization of the weights was chosen based on its AUPRC score on the test set. For figure 5A, all residues of a cAD 30mer were mutated to all other 19 amino acids and ADpred probability was computed. In Figure 5B the same approach was applied to cAD derivatives, and ADpred results were compared to experimental results (Tuttle et al., 2019).

To search for AD-regions in full protein sequences, we rolled a 30-residue long window over the entire sequences and assigned the score to the residue in the middle (the 16^th^ position in the 30mer). Ordered domains were obtained from HHpred (Zimmermann et al., 2018) and d2p2.pro webserver (Oates et al., 2013).

### Analysis of predicted AD function by RT qPCR

To experimentally test ADpred and to demonstrate that the model captures more than the amino acid composition of input sequences, we designed 30-mers with the same amino acid composition (and hence log-odds scores, Fig. 2A) but in the opposite extremes of the scale of ADpred probabilities. We picked sequences from low to high log-odds scores (A to E in **Fig 6A**) and permuted the order of amino acids in each of these sequences 10,000 times. We sorted each library of 10,000 sequences by their ADpred probabilities. We then selected peptides with high and low prediction scores and tested them for in vivo function by fusion to the Gcn4 DNA binding domain (**Supplementary Methods)**.

### Proteome analysis for ADs

To search for ADs in full length yeast protein sequences, a window of 30 residues was scanned along all annotated protein sequences in yeast (data from Saccharomyces Genome Database). and the ADpred probability for the window was assigned to the central amino acid in the window (the 16^th^). We calculated p-values for the enrichment of ADs in the set of transcription factors compared to the yeast proteome with the hypergeometric test as follows. The summed lengths *M* of all proteins in the proteome corresponds to the population size, and the summed length *N* of all transcription factor sequences corresponds to the “labeled” part of the population. Sites with five or more contiguous residues with a score ≥ 0.8 correspond to the samples drawn. Suppose there are *m* such sites, *n* of which are lying within transcription factors. The p-value for the hypergeometric test is the probability to obtain k or more sites within the transcription factors.

The enrichment is computed as

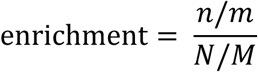

and the p-value corresponds to:

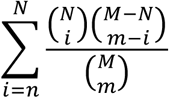

Implemented in scipy.stats.hypergeom as the survival function. Disorder and secondary structure predictions were calculated with PSIPRED 4.0.1(Cuff and Barton, 2000) and IUPred 1.0 (Dosztányi, 2017).

### 9aaAD motif were searched using *re* python library for regular expressions

The motifs searched are the most and less stringent (Prosite syntax): *[MDENQSTYG]{KRHCGP}[ILVFWM]{KRHCGP}{CGP}{KRHCGP}[ILVFWM] [ILVFWMAY]{KRHC}* and *[MDENQSTYCPGA]X[ILVFWMAY]{KRHCGP}{CGP}{CGP}[ILVFWMAY]XX*).

## Author Contributions

Jo_S and SH conceived the project, AE, LK, LW, Jo_ S and SH designed the experiments, AE, LW, and JF did the wet lab work, AE, LK, SSJ, Jo_S, and Ja_S performed computational analysis, AE, SH, Jo_S and SSJ wrote the manuscript, AE prepared figures, and all authors edited and approved the manuscript.

## Acknowledgements

We thank Steven Petesch for initial work on the project, Meera Hahn for advice on the deep learning NN, and Ziga Avsec for discussions on analysis of ADpred results. We also thank Kevin Struhl and Mark Ptashne for discussions and Brad Langhorst, and Lisa Tuttle for comments on the manuscript.

## Figure Legends

**Figure S1.**
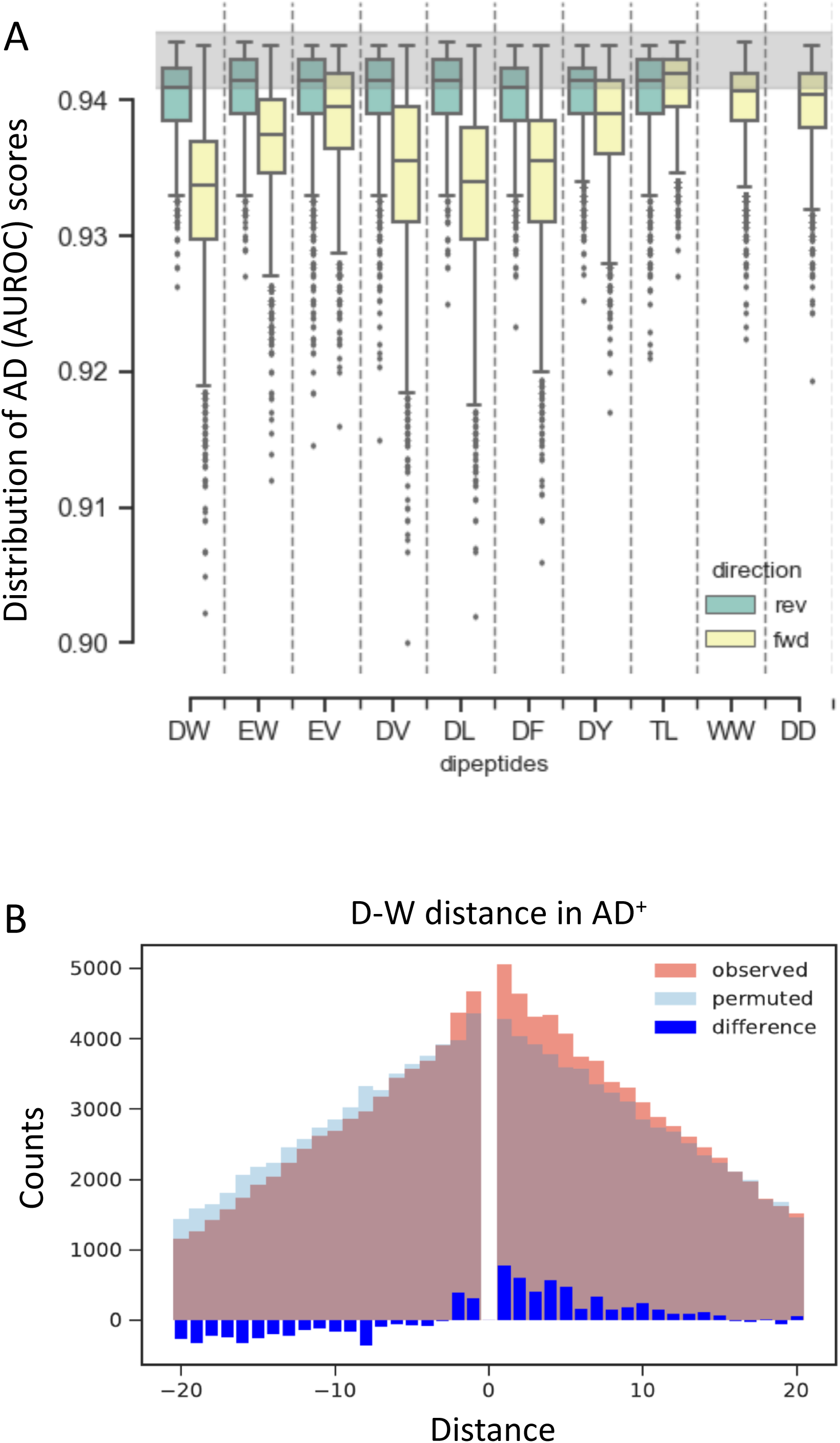
**A)** Distribution of AD scores obtained when a dipeptide coefficient is swapped to one of the 399 others (Methods and text). Reverse: WD instead of DW etc. **B)** Observed (red) and expected (blue) frequencies of DxW in the positive set depending on the length of x between 0 and +20 residues, and analogously for WxD plotted versus -x values. The expected frequencies are computed by randomly permuting the positive sequences. The difference between observed and expected frequencies is shown in dark blue.

**Figure S2.**
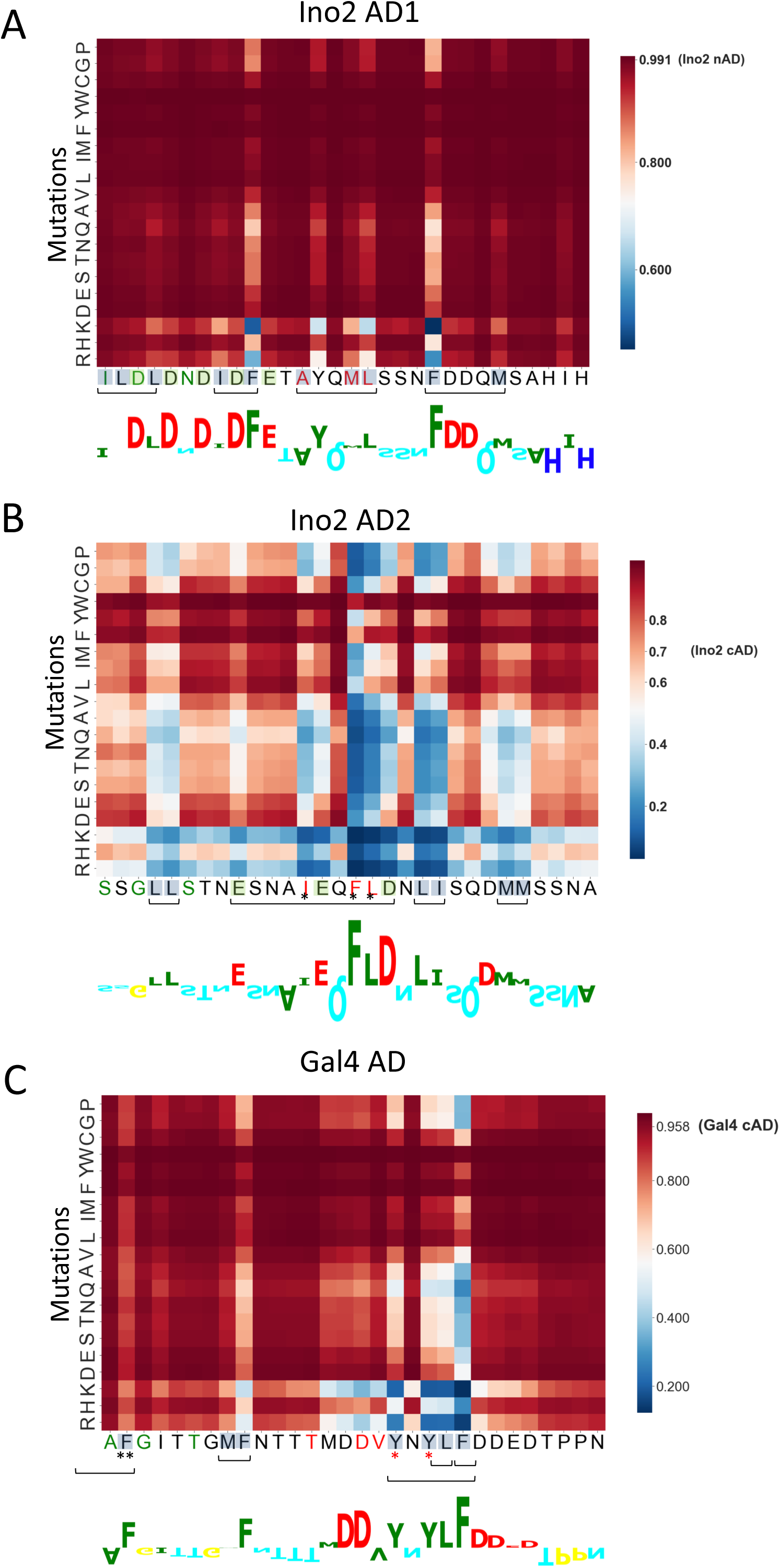
Prediction of important residues within yeast ADs and comparison with in vivo analysis. ADpred scores predicting the probability of AD function for all possible single amino acid mutations of the two Ino2 ADs and Gal4 AD. An increase in AD probability is darker red, a decrease blue. For comparison, results from an in vivo analysis where double or triple Ala substitutions were assayed for AD function (Pacheco et al., 2018; Tuttle et al., 2019). Conserved hydrophobic and acidic residues that were mutated are shown in blue and green, respectively. Double or triple Ala mutations resulting in less than ∼ 50% AD function are marked with brackets below the x-label. For Ino2 AD1, conserved residues that ADpred predicts to be important but not tested experimentally are indicated by: *. For Gal4 mutations, residue F849 (marked with **) was mutated in conjunction with Y846 and this derivative has 47% WT activity. Red asterisk marks Gal4 residues Y865 and Y867, which have ≥75% WT function when individually mutated to Ala.

**Figure S3.**
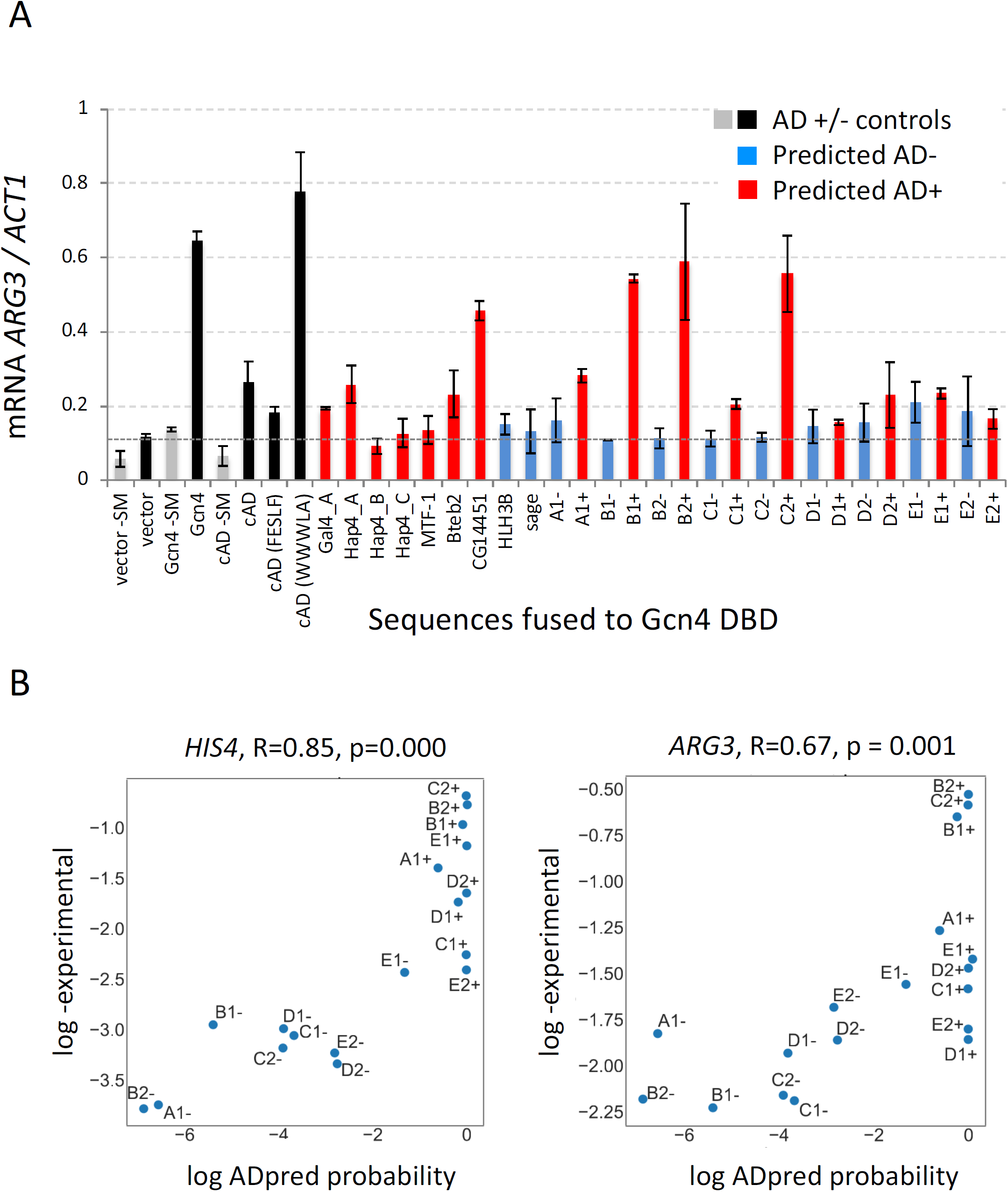
(**A**) Same as Fig. 6A but measuring AD function using RT qPCR quantitation of mRNA at yeast *ARG3*. Red bars = sequences with high ADpred probability; blue bars = low ADpred probability (See Fig S3B for ADpred scores). All samples were treated with SM unless otherwise indicated. Dotted horizontal line: level of SM-induced transcription in cells lacking Gcn4. (**B**) Scatter plot of the logarithm of ADpred probabilities versus log-experimental qPCR results obtained on *HIS4* and *ARG3* mRNAs. Pearson correlation and p-value for a two-sided hypothesis test (where the null hypothesis corresponds to slope=0) are indicated.

**Figure S4.**
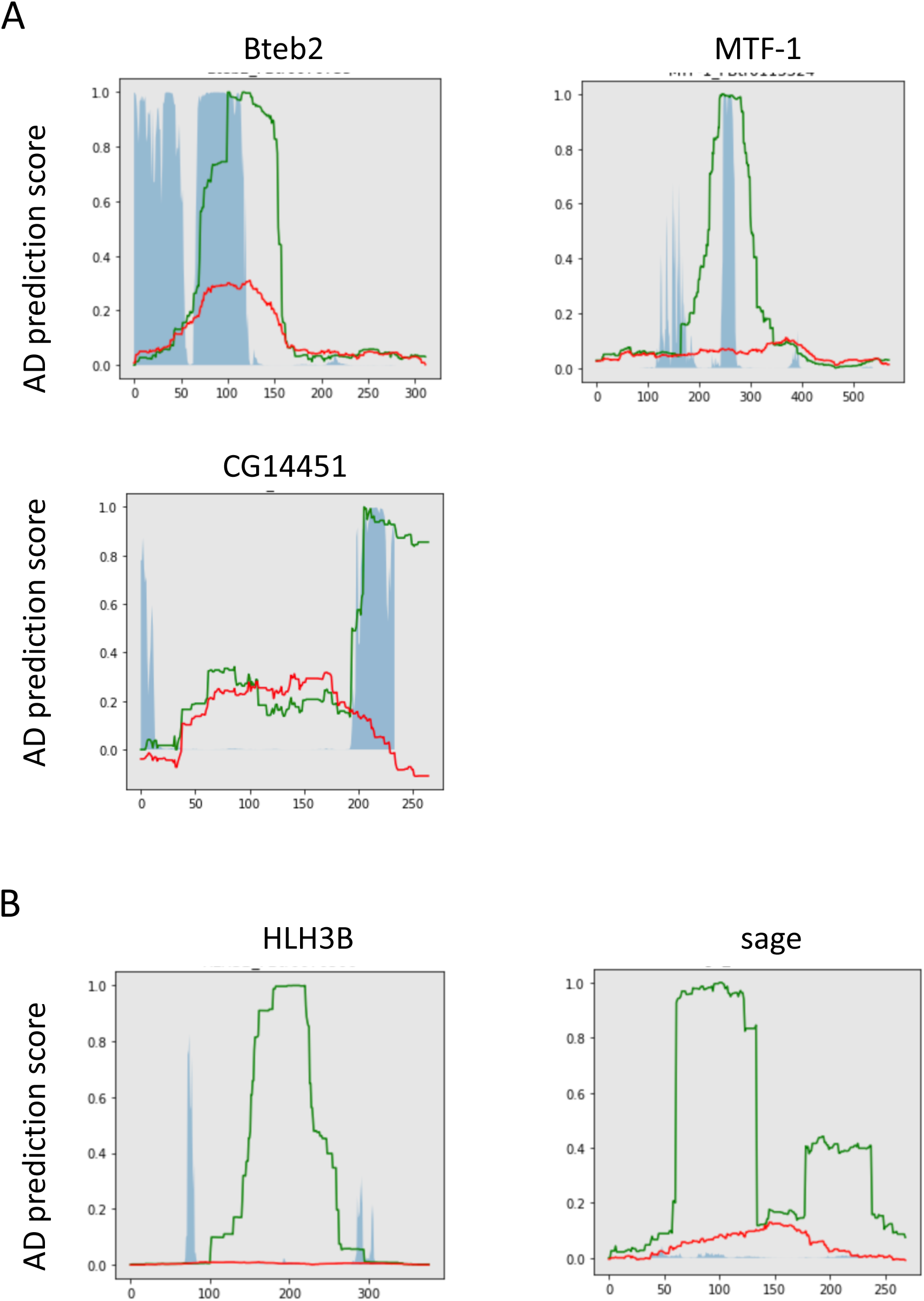
Five AD regions identified in a medium throughput screen in *Drosophila* (Arnold et al., 2018) that were used for functional assays in Fig 6A. Blue shading: ADpred scores; green lines: regions with AD function in *Drosophila*, red lines: background levels from *Drosophila* screen.

**Fig S5.**
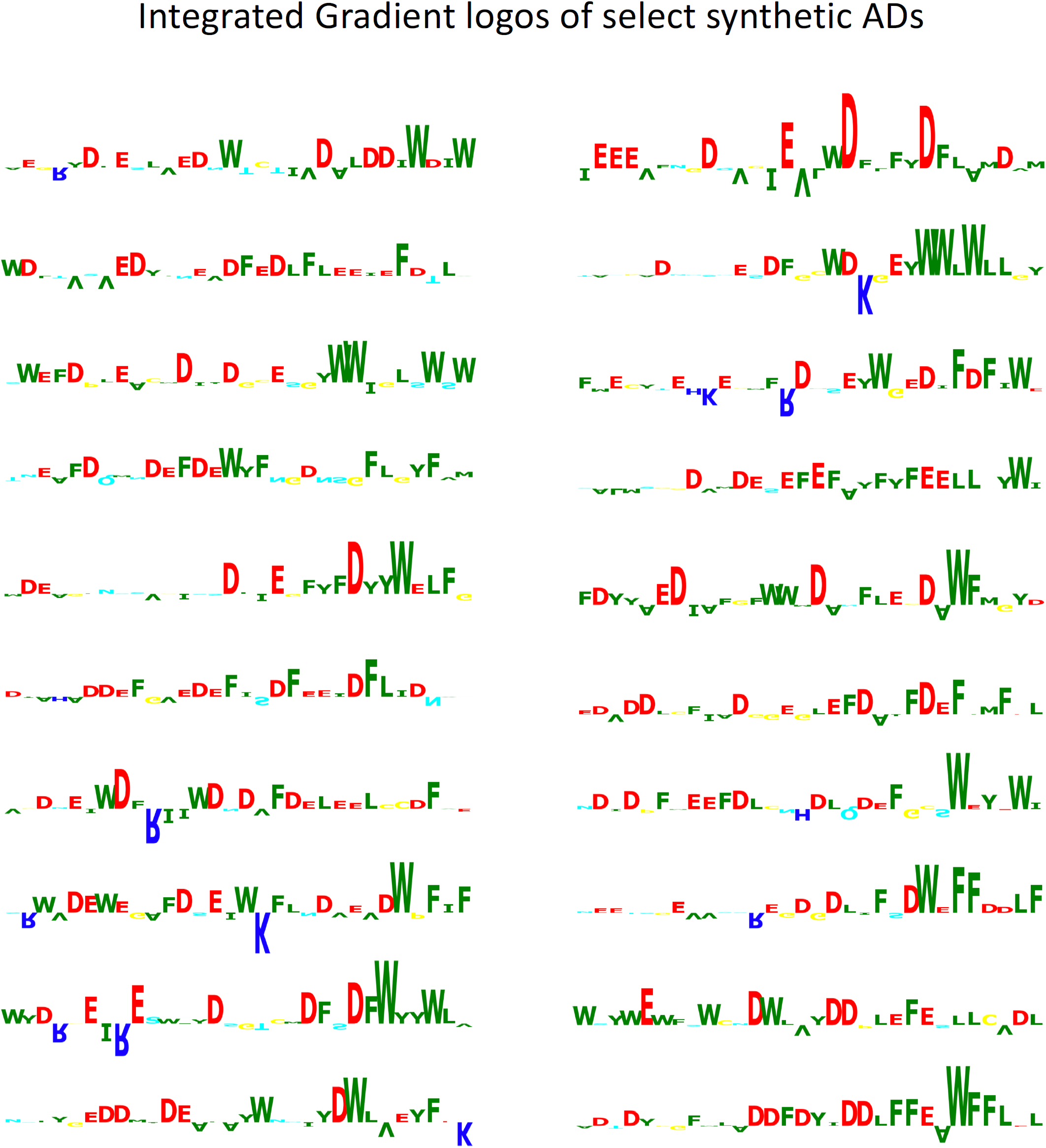
Shown are 20 of the top scoring synthetic ADs from the AD-positive set analyzed with Integrated Gradients. Logos are drawn as in Fig 6B.

**Figure S6.**
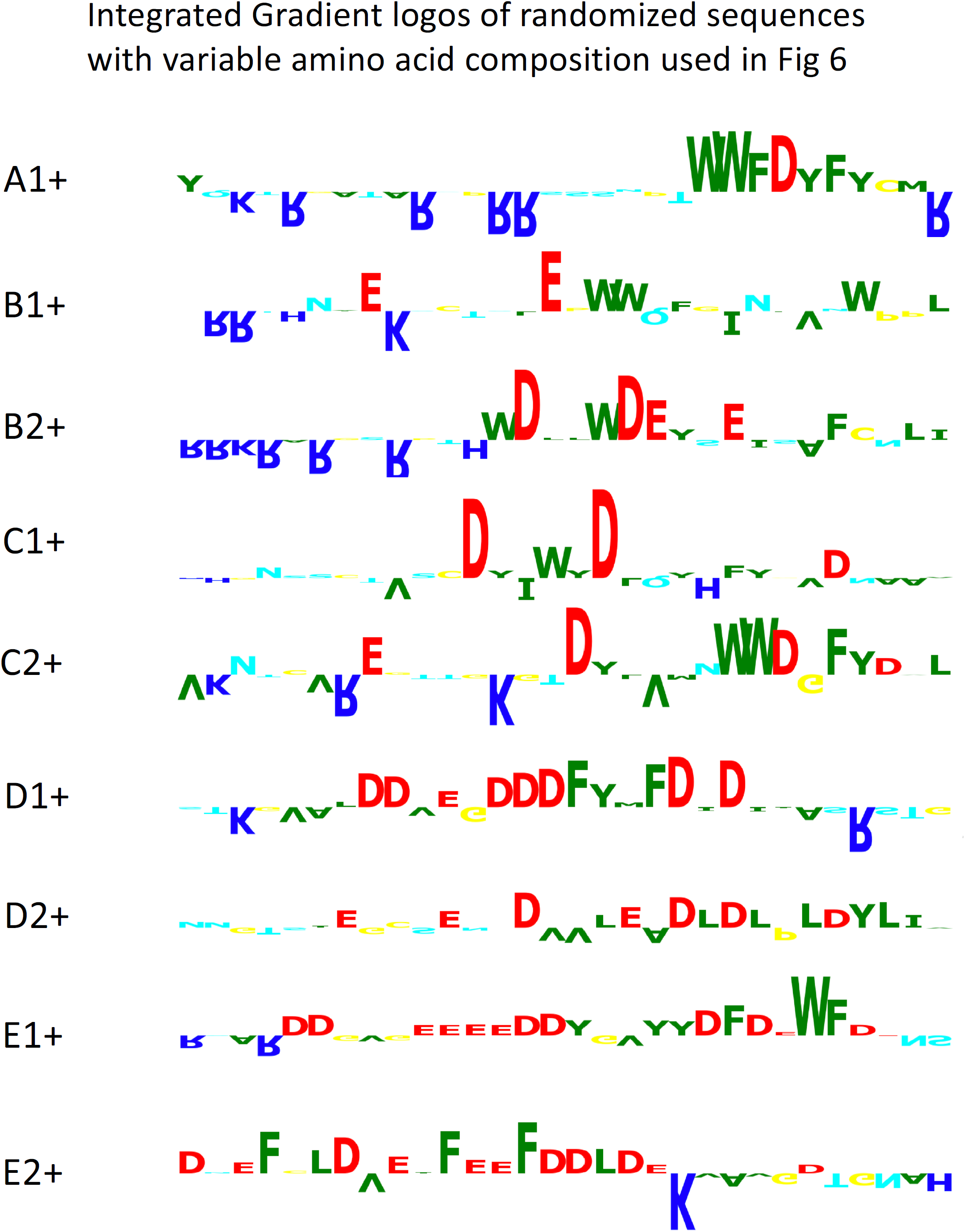
Shown is the Integrated Gradient analysis of predicted AD-positive peptides that were assayed for function in Fig 6A. Labeling of the peptides is as in Fig 6A and indicates amino acid composition with A being most unfavorable.

**Figure S7.**
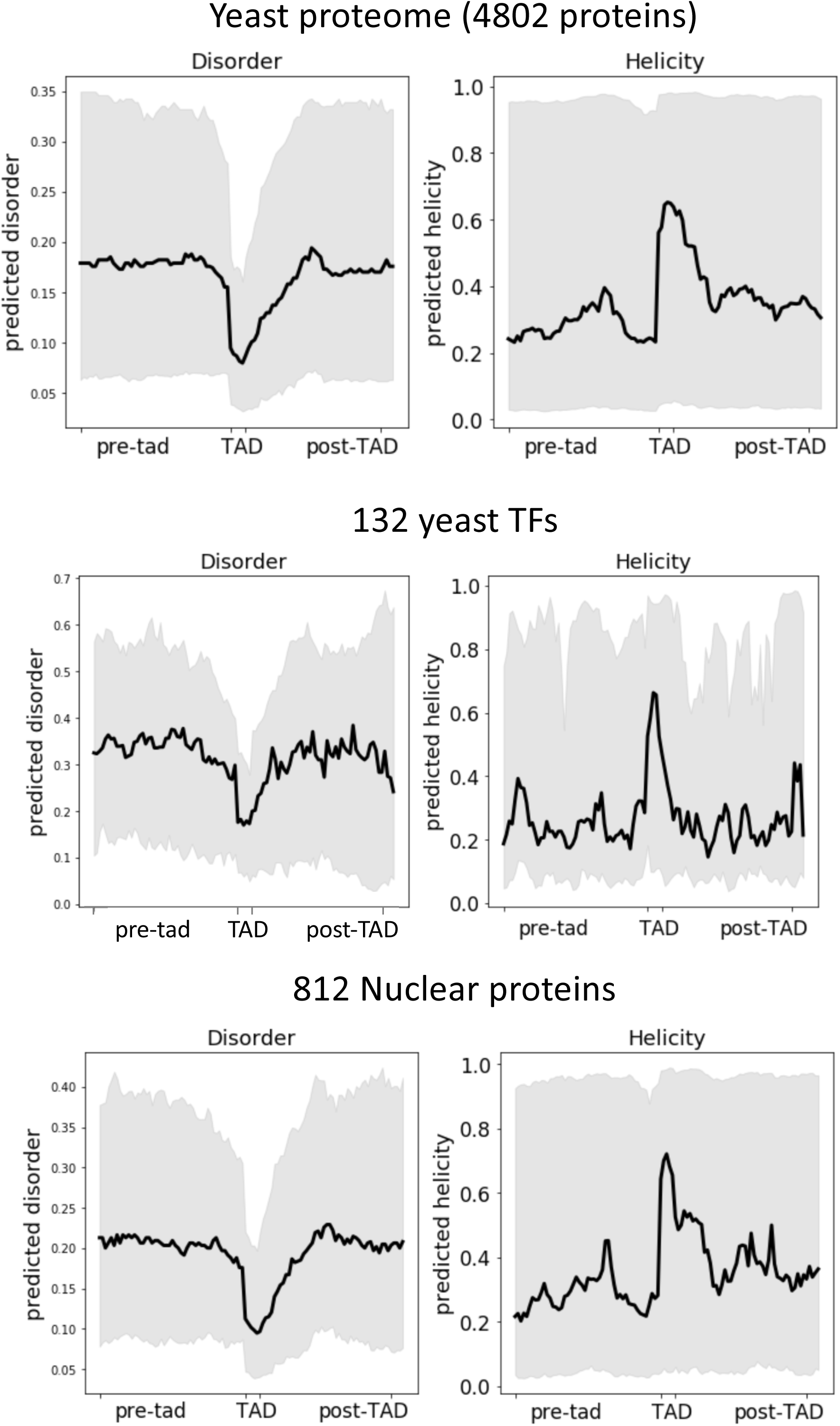
Structural properties of regions surrounding predicted ADs in yeast transcription factors. In the upper panel the whole proteome (only proteins with reported localization) is analyzed, a short list of 132 proteins considered transcription factors are analyzed in the middle panel (same as in Fig 7B) and 812 nuclear proteins are analyzed in the lower panel. “x” axis is composed of the 5 central residues of the predicted AD surrounded by 50 residues on each side. Black thick line represents the median and the grey shadowed area represents the 25^th^ to 75^th^ percentile.

## Supplementary Files and Tables

### Supplementary Methods

**Table S1** (open with text editor)

Unsorted list of AD-positive and AD-negative sequences with:

- List of AD-positive and negative sequences
- Distribution of sequences in the 4 bins
- Calculated AD-enrichment score

**Table S2**

- RT qPCR results
- Sequence of the synthetic ADs tested, and their DNA coding sequences

**Table S3**

- Set of 132 yeast TFs used in the AD search and predicted AD location for the longest AD in each factor.

**Table S4** (in Supplemental Methods)

Performance metrics of the regression and deep learning models.

